# Advanced models of lobular breast cancer metastasis capture clinical organ tropism, endocrine response, and bone remodeling

**DOI:** 10.64898/2026.03.13.711653

**Authors:** Joseph L. Sottnik, Mary E. Buchanan, Maria J Contreras-Zárate, Trinh C Pham, Maggie Musick, Thu H. Truong, Diana M Cittelly, Julie H. Ostrander, Matthew J. Sikora

## Abstract

Patients with invasive lobular carcinoma of the breast (ILC) are at high risk of long-term recurrence and metastatic progression with poor prognoses due to delayed detection and treatment-refractory disease. Unfortunately, few models are available to investigate metastatic ILC (mILC) and understand the unique metastatic patterns and phenotypes, including abdominal metastases, leptomeningeal disease, and mixed osteosclerotic/lytic bone metastases. Therefore, we expanded upon the previously established mammary intraductal (MIND) cell line xenograft model by supplementing mice with low-dose estradiol to promote disease progression. We observed spontaneous multi-organ spread from the mammary gland to common and mILC-specific tissues, with micro-metastatic disease as early as 12 weeks post-engraftment and macro-metastatic disease in 24-30 weeks, without the need for primary tumor resection. Primary and metastatic tumors remain highly endocrine responsive, allowing for the evaluation of novel therapeutics in the setting of disseminated metastasis. Derivative cell lines were isolated from various metastatic lesions, a total of 13 derivates from 7 sites across three hosts, and were found to have shared gene expression changes related to metabolism and intercellular signaling. Focusing on bone-derived variant cells as bone is the most common site for mILC to present, we found that bone-derived variant lines maintain multi-organ metastatic potential upon rechallenge by MIND or intratibial injection, despite increased aggressiveness and maintained endocrine response. Notably, bone lesions from either challenge route showed mixed osteosclerotic/lytic features characteristic to clinical ILC. Accordingly, we found that conditioned medium from ILC cells and the mILC bone-derived variants induce osteoblast differentiation and suppressed osteoclast differentiation *in vitro*, consistent with their effect on bone remodeling *in vivo* and in clinical disease. Together, the models developed herein can be utilized to understand the unique metastatic processes of mILC, and to investigate new therapeutic combinations in the setting of endocrine-responsive primary and metastatic ILC.

## INTRODUCTION

Invasive lobular carcinoma (ILC) of the breast accounts for ∼15% of new breast cancer diagnoses and affects >46,000 women in the US annually. ILC is primarily defined by its unique discohesive single-file infiltrative growth pattern, associated with loss of *CDH1*/E-cadherin and ∼95% estrogen receptor α (ER)-positivity. ILC is remarkable for increased risk of long-term recurrence, 5-8 years after initial diagnosis, with ≥30% increased risk of long-term recurrence relative to breast cancer of no special type (NST, aka ductal carcinoma) [1-5]. Management of ILC recurrence and metastatic ILC (mILC) is confounded by a unique pattern of metastasis compared to NST, with mILC spreading to the peritoneum, gastrointestinal tract (GI) and gynecologic tissues (uterus and ovaries) [6, 7]. These abdominal metastases are challenging to detect, resulting in delayed treatment and poor outcomes [8]. While the distinct metastatic tropism of ILC is presumed to be associated with E-cadherin loss, our understanding of the behavior of mILC is limited.

In addition to distinct metastatic tropism, the behavior of mILC at common sites of metastasis with NST, including brain and bone, is also distinct. NST metastases to the brain typically colonize the brain parenchyma, whereas mILC spreads to the leptomeninges, i.e. the structures surrounding and supporting the central nervous system [9]. While brain metastases are associated with poor outcomes, leptomeningeal disease (LMD) is associated with particularly poor outcomes [9-12]. LMD is typically undetected until rapid onset of neurological symptoms and is typically refractory to treatment, in part due to drug delivery challenges associated with the cerebral ventricular system [10]. Interestingly, mNST to the leptomeninges are associated with loss of E-cadherin, specifically within LMD and not synchronous metastases at other sites, suggesting that this hallmark of ILC facilitates spread to the leptomeninges [10, 12].

Bone is the most common site of ILC metastasis, and often the first site diagnosed [13]. 80% of patients with mILC will develop bone metastases, and those with bone-only disease have poorer outcomes compared to mNST patients [7, 8, 13]. Numerous case reports describe mILC bone metastases as diffuse with heterogeneous osteosclerosis [14-19], in stark contrast to the predominantly osteolytic behavior of mNST [20]. It is unknown why mILC lesions are often osteosclerotic, and the extent/frequency of sclerotic lesions has not been robustly examined, but this complicates diagnosis of advanced disease. Patients with *de novo* mILC in bone may be misdiagnosed with sclerotic diseases like osteopoikilosis and tuberous sclerosis [17]. Despite the heterogenous behavior of mILC in bone, NST and ILC bone metastases are all treated with anti-resorptive therapies such as bisphosphonates (eg zoledronic acid) or anti-RANKL (Denosumab) therapies [21, 22].

New models to study the unique phenotype of mILC are necessary to understand mILC tropism and its distinct behavior within the metastatic niche, toward developing new strategies for detection, monitoring, and treatment. However, use of ILC cell lines in common xenografting models has been limited to date. Recent use of mammary intraductal (MIND) engraftment showed that ILC cell lines MDA-MB-134VI (MM134) and SUM44PE (44PE) spontaneously spread from the mammary gland to common and ILC-specific sites [23]. In the original study, metastases required ≥6 months to be detectable at distant sites by *ex vivo* bioluminescence, and the majority of spread remained micro-metastatic even 9-12 months after engraftment, limiting the potential of MIND for modeling ILC metastasis. Importantly, a key feature of these original studies with the MIND model was the omission of exogenous estrogen, toward modeling low estrogen levels in post-menopausal women or those treated with aromatase inhibitors. However, we considered that given the remarkable estrogen-dependence of ILC cells *in vitro* [24] and *in vivo* [24, 25], estrogens produced by the mouse ovary may be simply insufficient. Recent work with patient-derived xenografts (PDX) of breast cancer also identified alternatives for estrogen delivery (over commercially available sub-cutaneous pellets) that better mimic human physiological levels of estrogen [26]. Based on these observations, we revisited MIND engraftment of ILC cells *in vivo*. Using the MIND model, we were able to recapitulate ILC-specific mILC tropism that ultimately presented as macro-metastatic and lethal mILC. From these models, we further explored mILC tropism and behavior, focusing on the bone microenvironment, wherein MIND mILC bone lesions closely mimic the clinical heterogeneity of mILC in bone. These models provide a new window to understand the behavior of mILC in the metastatic niche.

## MATERIALS AND METHODS

### Cell Culture

MDA-MB-134VI (MM134; ATCC HTB-23) and SUM44PE (44PE; Asterand/BioIVT) were maintained as previously described [27]. HCC1428 (ATCC CRL-2327) and T47D (Sartorius Lab, University of Colorado) were maintained as previously described [27]. UCD4-luc-GFP (Sartorius Lab, University of Colorado) were maintained as previously described [28]. 293FT (Invitrogen) were maintained as directed. CAMA-1 (ATCC HTB-21), UACC-3133 (CRL-2988), and ZR-75-30 (CRL-1504) were maintained as described [29]. MC3T3-E1 (ATCC CRL-2593) and Raw264.7 (ATCC TIB-71) were maintained as described [30, 31]. Cells were incubated at standard conditions (37° C and 5% CO_2_). All cell lines were regularly tested and confirmed mycoplasma-negative using a MycoAlert kit (Lonza), and authenticated by STR profiling at the U. Colorado Anschutz Cell Technologies Shared Resource (RRID: SCR_021982). Estradiol (E2) was purchased from Sigma-Aldrich and diluted in ethanol.

### Development of luciferase stable expressing cell lines

pLenti CMV V5-LUC Blast (w567-1; Addgene #21474, a gift from Eric Campeau). Lentiviral particles were assembled in 293FT cells using psPAX2 (Addgene #12260, a gift from Didier Trono) and pMD2.G (Addgene #12259, a gift from Didier Trono) via transfection. Lentiviral transduction of MM134, 44PE, T47D, and HCC1428 was followed by blasticidin selection of bulk cells to maintain population heterogeneity and potential metastatic phenotype. Transduced cell lines are defined by ‘-luc’ throughout. Luciferase expression was confirmed using a Biotek Neo2 plate reader.

### Cell proliferation and doubling time

Double-stranded DNA was quantified using Hoechst 33258 fluorescence as previously described and used as a surrogate for cell number [27]. MM134-luc (1×10^4^ cells per well) and 44PE-luc (1×10^4^ cells per well), including 44PE-luc derivative lines, were plated in a 96-well format. 24 hours after plating, cells were treated with the noted concentrations of estradiol (E2) and incubated under standard conditions for up to 6 days. In addition, cells were hormone-deprived in medium supplemented with charcoal-stripped FBS (CSS) as previously described, then treated with vehicle (0.01% ethanol) or 100pM estradiol for the duration of the assay [27, 32].

### Cell cycle analysis

Cell cycle analysis was performed by plating cells for at least 72 hours, prior to staining [27, 33]. Cells were trypsinized, fixed (70% ice cold ethanol for 30 minutes), and stained in PI solution. Cells were analyzed on a Gallios 561 cytometer (Beckman-Coulter) and analyzed in Kaluza v2.1 (Beckman-Coulter).

### RNA-seq analysis

For RNA-seq of derivative cell lines, cells were plated at 4x10^5^ cells per well under standard conditions and incubated for 72 hours. Cells were washed and RNA extracted using the illustra RNAspin Mini Kit (Cytiva) according to manufacturer instructions. RNA was submitted to Novogene for library preparation and next generation sequencing. Data was prepared for analysis by the University of Colorado Cancer Center Bioinformatics and Biostatistics Shared Resource Core and analyzed using the cores ‘RNA-seq Analysis Tool.” For +/-E2 RNA-seq, cells were suspended in DNA/RNA Shield (Zymo Research) and submitted to Plasmidsaurus for library creation and sequencing. Data analysis was as described above.

### Osteoblast and osteoclast differentiation assays

MC3T3-E1 (osteoblast) and Raw264.7 (osteoclast) differentiation assays were performed using previously published STAR methods [34]. Conditioned media (CM) was prepared by plating tumor cells at 75% confluency and changing media to 0.1% FBS in DMEM/F12 media after 24 hours. After 5 days, CM was removed from the donor cells and filtered using a 0.45µm steriflip (Millipore Sigma). Total protein was normalized by diluting with additional DMEM/F12.

For osteoblast (OB) differentiation, 5x10^4^ MC3T3-E1 cells were plated in 24-well plates. 24 hours after plating, media was changed to 50:50 MC3T3-E1:CM and supplemented with 50µg/ml ascorbic acid (Sigma-Aldrich), 10mM β-glycerophosphate (Sigma-Aldrich), and 5mM CaCl_2_ (Sigma-Aldrich) [35]. Media was refreshed every 3 days for the duration of the assay. After 28 days, alizarin red staining was performed by first fixing cells in 4% paraformaldehyde for 2 hours at room temperature. Plates were washed with water and stained with 2% Alizarin Red S (Sigma-Aldrich) pH4.2 for 1 hour at room temperature. Stain was discarded and plates washed and dried. Plates were imaged with an Olympus APX-100. Stain was eluted with 10% acetic acid and gentle agitation for 1 hour at room temperature. Eluted stain was transferred to a 96-well plate (in triplicate) to measure optical absorbance at 405nM to determine the amount of mineralized calcium present.

For osteoclast (OC) differentiation, 2.5x10^4^ Raw264.7 cells were plated per well in a 24-well format. 24 hours after plating, media was changed to 50:50 Raw264.7:CM and supplemented with 50ng/ml of RANKL (PePro Tech). Media was refreshed after 48 hours. 4 days after CM initiation, Raw264.7 cells were fixed with 4% formaldehyde for 10 minutes at room temperature and then washed with absolute methanol. Cells were stained with May-Grunwald (Sigma-Aldrich) for 1 minute, washed, and then stained with Giemsa-Azur solution (Sigma-Aldrich) and then washed with water and dried. Plates were imaged with an Olympus APX-100 microscope. Mature Osteoclast (mOC; >3 nuclei per cell) were manually counted in 10 regions of interest (ROI).

### Animals

All animal studies were performed in an AAALAC-approved facility with approval of the University of Colorado Anschutz Institutional Animal Care and Use Committee (IACUC). 6-8 week female NSG (NOD.Cg-*Prkdc^scid^ Il2rg^tm1Wjl^*/SzJ; Jackson Laboratory strain 005557) mice were used for all experiments. 5-10 mice per group were used for all *in vivo* experiments. One week prior to tumor challenge, mice were initiated on 30µM 17β-estradiol (E2) supplemented in the drinking water. Mice were allowed to drink and eat ad libitum for the duration of the study as previously described [33]. Estrogen withdrawal (EWD) was achieved by removing supplemental E2 and returning mice to standard water. Mice were weighed weekly as weight loss >10% and physical signs of stress were used as indicators of acute toxicity and metastatic progression. All mice were administered Buprenorphine-SR/ER (Wedgewood Pharmacy; Swedesboro, NJ 08085) for analgesia at the time of tumor challenge.

### Tumor challenge

For mammary intraductal (MIND) challenge, tumor cells were prepared for injection and diluted in HBSS. Mice were anesthetized with isoflurane, abdomen shaved, and challenged with 1×10^5^ cells, per site, in the bilateral #3 and #4 mammary fat pads as previously described [23]. For intracardiac (IC) injection, tumor cells were prepared as above. Mice were anesthetized, placed in dorsal recumbency, and the chest swabbed with ethanol, and challenged with 1×10^6^ cells via cardiac puncture as described [36]. For intratibial (IT) challenge, tumor cells were prepared as above. Mice were anesthetized, hind-limbs shaved and disinfected with ethanol. A 22g needle was used to drill a pilot hole into the tibial plateau and extended into the medullary cavity of the tibia. 1×10^6^ cells were injected into the intramedullary as described [37, 38].

### Xenogen imaging

Tumors were monitored by bioluminescent imaging (BLI) using an IVIS spectrum (Revvity) and Living Image 4.8.2 software (Revvity). Whole body images were taken every 2-4 weeks during tumor progression. At experiment conclusion, *ex-vivo* (post-mortem) BLI was accomplished immediately (within 15 minutes) after whole body imaging to determine metastatic burden in each organ. Total metastatic burden is the sum of all plates used for analysis. Background BLI signal for metastatic burden was taken from sham challenged mice undergoing the same imaging protocol as above.

### microCT

microCT analysis was performed by the University of Colorado Cancer Center Animal Imaging Irradiation Shared Resource Core Facility (RRID:SCR_021980) on neutral buffered formalin fixed hind limbs. Image analysis was performed using a PerkinElmer Quantum GX2 microCT scanner. DICOM data was visualized using MicroDicom (https://www.microdicom.com/), reconstructions performed with 3D Slicer (https://www.slicer.org/; [39]) and Analyze 15.0 (AnalyzeDirect, Inc).

### Histology

Histologic staining was performed by University of Colorado Cancer Center Pathology Shared Resource. Hematoxylin & Eosin (H&E), ER, and Ki67 were stained on formalin fixed paraffin embedded (FFPE) tissues. For bone, after microCT analysis, bones were decalcified in 14% EDTA for 2 weeks and fixed in 10% neutral buffered formalin overnight before paraffin embedding.

### Development of metastatic lesion-derived cell line variants

For soft tissues, BLI+ disease identified on *ex vivo* analysis was excised and digested as previously described [40]. Briefly, tumors were dissociated and digested in Accumax (STEMCELL Technologies) for 1 hour at 37°C. Remaining tissue fragments were pushed through a 70µm filter. Red blood cells (RBC) were lysed using ACK media (154.4 mM ammonium chloride, 10mM potassium bicarbonate, and 97.3uM EDTA) and washed prior to plating in cell line specific media. For bone metastases, tumors were isolated in a manner similar to osteoblasts [34]. Mechanically dissociated bone fragments were digested with Collagenase/Hyaluronidase (STEMCELL Technologies) for 1 hr with agitation at 37°C. DNAse and RNAse (Millipore Sigma) were added to the solution and incubated for 15 minutes at 37°C. RBC were lysed with ACK as above, washed, and cultured in cell lines specific media. All cultures were supplemented with antibiotic-antimycotic (Fisher Scientific) for two weeks after isolation. After expansion BLI intensity was confirmed using a Biotek Neo2 plate reader. Supplemental Table 1 describes the organotrophic derivative cell lines.

### Software

Statistical analyses and associated graphs were prepared with GraphPad Prism v10 (Dotmatics, Boston, MA, USA).

## RESULTS

### Modeling mILC in an estrogen-replete setting yields lethal macro-metastatic disease mirroring clinical mILC

Using the MIND engraftment model in NSG mice, we examined the impact of an estrogen-replete environment (ovarian intact mice supplemented with 30µM E2 in the drinking water [26, 41]) on the progression of ILC xenografts in the murine mammary gland. Mice were challenged by MIND with ER+ ILC models MM134, 44PE, or MM330. After 8-20 weeks, mammary glands were harvested to examine lesion growth and histology, and to evaluate the impact of exogenous E2 on tumor progression. Consistent with the estrogen-responsiveness of these lines [24, 42], supplemental E2 increased the number, size, and invasiveness of lesions for all three ILC models (**Figure 1A/B**). Absent exogenous E2, lesions were small and contained within the mammary duct, with little to no invasion into local stroma/adipose tissue, but nearly all lesions in E2-supplemented mice showed clinical ILC-like invasion, e.g. invasion between and around adipose cells.

**Figure 1:**
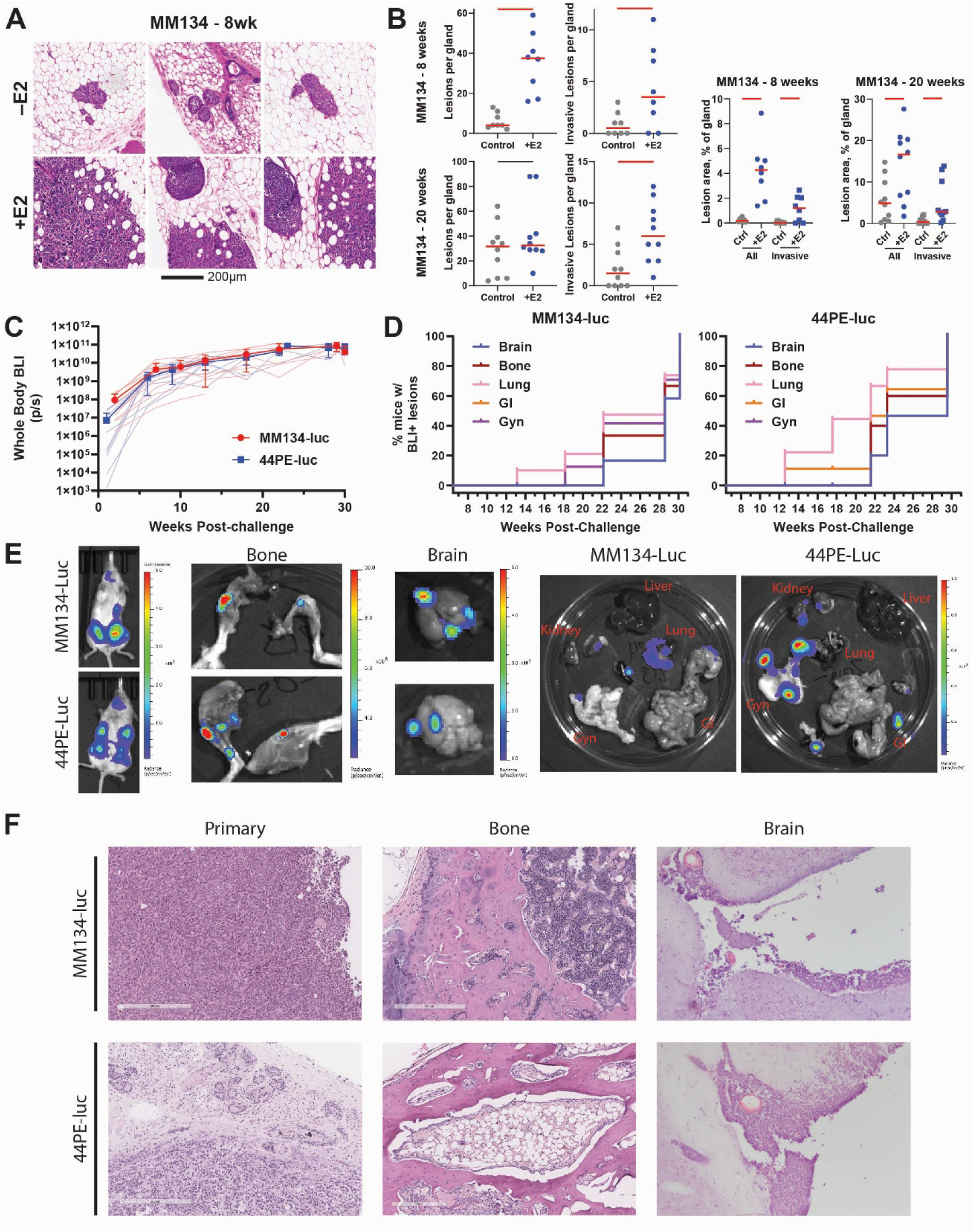
Development of mILC models with estrogen supplementation. A) Representative histologic comparison of MIND tumors grown without (-E2) and with (+E2) supplemental estrogen in the drinking water 8 weeks after challenge. B) Quantification of histologic for number of lesions, number of invasive lesions, and tumor area compared by t-test; red bars p<0.05. C) Whole body tumor burden as measured by BLI for MM134-luc and 44PE-luc MIND challenged mice (n=10 per line) supplemented with 30µM estrogen in the drinking water. D) Serially sacked cohorts of mice to identify metastatic progression in distal organs. E) Representative images of MM134-luc and 44PE-luc mice at the time of sacrifice and *ex vivo* BLI analysis of bone (hind-limbs), brain, and visceral organs. F) Representative H&E images of MM134-luc and 44PE-luc primary tumor and bone metastases.

Based on these observations in primary lesions, we transduced a luciferase reporter into MM134 and 44PE cells (referred to as MM134-luc and 44PE-luc) to further track tumor growth and metastatic progression with estrogen supplementation. To define metastatic progression, mice were euthanized in cohorts over the course of 30 weeks to perform bioluminescence imaging (BLI) *ex vivo* (**Figure 1C/D**); *ex vivo* BLI allows for detection of metastatic disease that is obscured in whole body imaging. Throughout the time course, primary tumors were contained within the mammary fat pad and did not grow to a size requiring resection or meeting humane endpoints.

Micro-metastatic development (i.e. BLI signal detection in organs *ex vivo*) was observed as early as 12 weeks in both models, and by 30 weeks post-challenge all mice had multi-organ metastatic disease to the gynecologic tissues (Gyn; uterus and ovaries), kidney/adrenal gland, lung, brain, gastrointestinal (GI) tract, and bone (**Figure 1E**), consistent with common and ILC-specific sites observed clinically [7]. We did not detect liver metastases in either ILC model, mirroring the reduced frequency of mILC in the liver relative to mNST [7]. Importantly, while we did not observe progression beyond micro-metastatic disease in the lung, lesions across other sites progressed to macro-metastatic, symptomatic disease during the time course.

Metastases to the ovary and uterus were typically large and palpable 16 weeks post-challenge in both models, and histology showed metastases within the ovarian parenchyma (**Supplemental Figure 1**). Strikingly, at ≥24 weeks, mice presented with central nervous system-related symptoms (logrolling, rotating, facial grimace/paralysis) which could be attributed to brain metastases and necessitated humane euthanasia in >40% of both MM134-luc and 44PE-luc animals. Histologic analysis of brain showed that metastatic cells were present in and largely confined to the leptomeninges, mirroring clinical LMD with mILC [9, 10], with some invasion into the brain parenchyma in 44PE-luc (**Figure 1F** and **Supplemental Figure 1**). BLI+ Bone metastases were readily detectable in the hind-limbs and confirmed by histology (**Figure 1F**). IHC analysis of Ki67 showed metastatic lesion proliferation comparable to the primary tumors (**Figure 1F**). Notably, metastases retained ER expression in all metastatic tissues (**Figure 1F and Supplemental Figure 1**), suggesting that metastases may maintain E2-sensitivity.

We characterized additional ER+ ILC-like models [43] CAMA1 (luminal-like), UACC-3133 (*ERBB2*-mutant), and ZR-75-30 (HER2-positive) with luciferase-expressing cells, engrafting cells by MIND in E2-supplemented mice to track tumor growth and metastatic progression. These ILC-like models grew similarly to MM134 and 44PE with metastatic progression to multiple tissues, including bone, brain, lung, and ILC-specific sites within 23 weeks of challenge (**Supplemental Figure 2**).

These data support that supplementation of MIND xenografts of ILC cell lines with low-dose E2 advances the rate of growth and metastatic progression, facilitating the development of macro-metastatic disease that mirrors the presentation of clinical mILC. Notably, we did not observe any E2-related toxicity in mice, including uterine enlargement or urogenital infections, consistent with much lower E2 levels than other delivery methods [26, 44, 45]. Thus, exogenous E2 may make the study of ILC spontaneous metastases from the mammary gland tractable, both for metastatic mechanisms and for modeling treatment response.

### MIND mILC can model response to standard-of-care endocrine therapy and novel therapeutics

We next tested whether estrogen withdrawal (removal of E2 from drinking water) after the establishment of MIND metastases impacted the growth of primary and metastatic lesions, to mimic progression in an E2-repressed setting. Intact mice have circulating levels of estradiol (2.7± 1 pg/ml) similar to those of post-menopausal women (1.3-4.9 pg/ml) [46] whereas the removal of estrogen from the drinking water recapitulates aromatase inhibition (AI; a significant, though incomplete, drop in circulating estrogen levels) in patients. Since we recently reported on a novel form of PARP inhibitor sensitivity in ILC, including against orthotopic cell line xenografts and patient-derived xenografts [25], we also included the PARP inhibitor (PARPi) talazoparib in combination with estrogen withdrawal. In these treatment studies, MIND xenografts of MM134-luc and 44PE-luc were allowed to establish for 15 weeks to ensure metastatic engraftment prior to initiating treatment. As above, we assessed whole body BLI and *ex vivo* BLI to measure metastatic burden.

E2 withdrawal significantly reduced tumor growth (whole body BLI) in both MM134-luc and 44PE-luc (**Figure 2A-B**), consistent with the endocrine responsiveness of these models. Control tumors (with E2) led to symptomatic metastatic disease requiring euthanasia by ∼25 weeks or ∼30 weeks in MM134 or 44PE, respectively, and *ex vivo* BLI confirmed extensive multi-organ metastases at these endpoints. Estrogen withdrawal (labeled as ‘AI’ for modeling aromatase inhibitors) slowed metastatic progression, as most mice remained asymptomatic out to ∼37 weeks and ∼33 weeks in MM134 or 44PE, respectively (**Figure 2C-D**). Both models eventually had age-related health issues in the AI-treated groups, as mice reached ∼1 year old, but not metastasis-related symptoms seen in control mice. *Ex vivo* BLI confirmed that metastatic burden was reduced by E2 withdrawal, and time to lethal levels of metastatic burden were extended in both models (**Figure 2E-F**).

**Figure 2:**
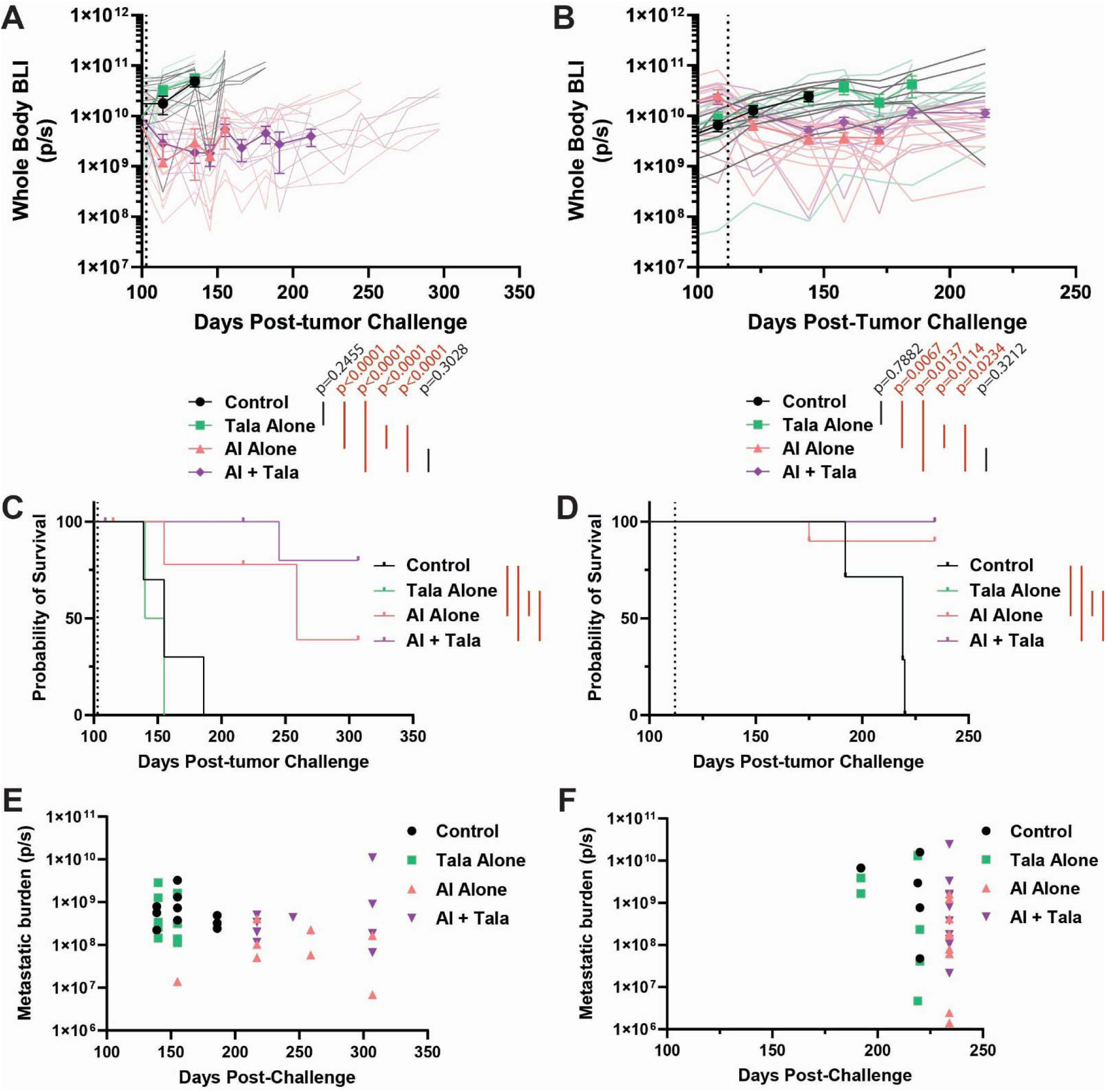
Therapeutic evaluation of combined estrogen withdrawal and talazoparib treatment in MIND models. MM134-luc (A) and 44PE-luc (B) tumors challenged by MIND (n=10/group) and allowed to establish prior to treatment initiation of estrogen withdrawal (AI; mimicking aromatase inhibition), 0.33 mg/kg talazoparib (Tala alone), or a combination of estrogen withdrawal with talazoparib treatment (AI+Tala). A) MM134 and B) 44PE whole body BLI after treatment initiation (vertical dotted line). Darker lines represent group averages until euthanasia of first mouse per group. Faded lines represent individual mice until their individual time of euthanasia. Red brackets show statistically significant (p<0.05) comparisons by repeated measures two-way ANOVA treatment values. C) MM134-luc and D) 44PE-luc overall survival. Red brackets show statistically significant comparisons (p<0.05) using pairwise survival analysis. Total metastatic burden depicted by *ex vivo* BLI compared to the time of euthanasia in MM134-luc (E) and 44PE-luc (F).

As we observed in orthotopic models [25], PARPi alone did not impact tumor growth, but combined AI+PARPi further slowed primary and metastatic tumor growth and extended survival (though mice ultimately had similar age-related health issues as the AI alone arms). These data support that ILC metastases via MIND are endocrine responsive and can be used to model the impact of treatment on metastatic progression.

### Gene expression analysis of ILC cell line derivatives from metastatic lesions suggests shared remodeling

To investigate the organotrophic phenotype of ILC metastases, BLI+ macro-metastases (**Figure 1**) were isolated and expanded *ex vivo* as a series of cell line variants. This series is derived from three MM134-luc host mice (n=4 derivative lines) and three 44PE-luc host mice (n=13 derivative lines), summarized in **Table 1**. After outgrowth, we confirmed cell line identity by *in vitro* luciferase positivity; we then used RNA-seq to compare 44PE derivatives to their parental line.

Differential gene expression analyses showed that despite being derived from independent mice, lesions, and sites, overall gene expression changes were remarkably consistent across the metastasis-derived variants (**Figure 3A**), and hundreds of genes were coordinately induced or repressed across sites (**Figure 3B**, **Supplemental File 1**). Upregulated pathways prominently feature lipid and phosphatidylinositol metabolism pathways (**Figure 3C**), while the MSigDB Hallmarks for epithelial-mesenchymal transition (EMT) and apical junction were downregulated (**Figure 3D**). Consistent with suppression of EMT signatures, signatures for mammary luminal epithelium were upregulated while signatures for mammary basal epithelium were downregulated (**Figure 3E**). Additionally, while the Hallmark for estrogen response early was enriched in downregulated genes (adj.p = 0.02 among genes downregulated in variants from at least 6 sites), changes in ER target gene expression (using the Estrogene signatures [47]) suggested a remodeling of ER target gene control rather than outright decreased ER activity (**Figure 3F**). Collectively, gene expression changes in metastasis-derived cell line variants suggest remodeling of cell metabolism and cell surface/adhesion properties are part of the metastatic cascade for ILC.

**Figure 3:**
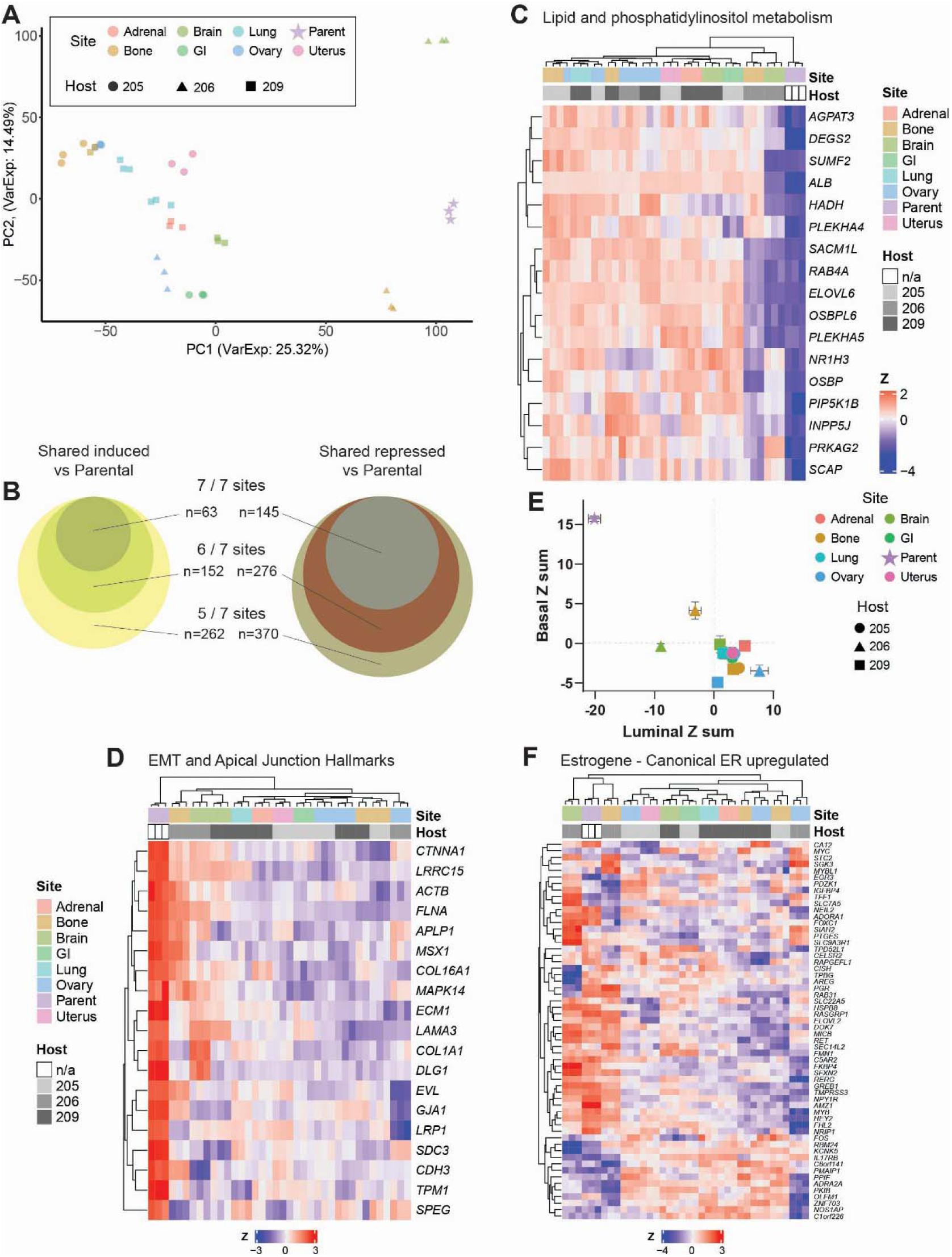
Derivation of organotrophic ILC metastases share a common transcriptomic phenotype. Spontaneous metastases from MIND challenged mice (Figure 1) were isolated and expanded *in vitro*. A) RNA-seq of the resulting organotrophic lines, isolated from 3 mice (numbers 205, 206, and 209) was performed and compared by PCA. B) Representation of induced and repressed genes shared by >5 metastatic sites compared to the parental cell line. Heatmap, and unbiased hierarchical clustering, of genes associated with lipid and phosphoinositol metabolism (C) or EMT and Apical Junction Hallmarks (D) of derivative cell lines compared to the parental line (Host = n/a). E) Basal vs Luminal phenotype z-score sums were calculated and assigned to derivative cell lines showing enrichment of the luminal phenotype in metastatic derivatives. F) Comparison of organotropic derivatives using canonical ER upregulated genes from Estrogene.

### Bone-derived ILC lines maintain the ability to metastasize to other organs

We focused on bone-derived 44PE variants to assess organ tropism, since bone is the most common site for mILC and typically the first site where mILC presents [13]. We challenged mice with 44PE-luc and bone-derived variants from two different hosts, 44PE-Bo206 and 44PE-Bo209, by three different approaches (all with E2 supplementation): 1) MIND engraftment, modeling metastatic dissemination from the primary tumor; 2) intracardiac injection (IC), modeling cell survival in circulation, extravasation, and colonization; 3) intratibial injection (IT), modeling growth directly in the bone microenvironment. In all models, growth was tracked by whole body BLI and metastatic burden measured by *ex vivo* BLI as above.

In all three xenograft models, the bone-derived variants grew significantly faster than the 44PE-luc parental cell line based on whole body BLI (**Figure 4A**). In MIND at 18 weeks post-engraftment, bone variants had ∼10-fold greater whole-body BLI than the parental 44PE (**Figure 4B**), while based on our prior studies with 44PE-luc (**Figures 1 and 2**) the parental cells would not reach a similar size until 30-32 weeks (**Supplemental Figure 3A**). We confirmed that increased tumor burden was not an artifact of luciferase expression as the bone-derived lines in fact had decreased luminosity with equivalent cell number *in vitro* compared to the parental line (**Supplemental Figure 3B**). Increased primary tumor growth in MIND corresponded with greater total metastatic burden with the bone derivatives (**Figure 4C**). However, while burden at other individual sites was accordingly increased for the bone derivatives, this was not the case for metastasis to the bone, which had equivalent BLI in parental and bone-derived cells (**Figure 4D**). By IC injection, a modest increase in whole body BLI was observed in the bone-derived models (**Figure 4E**) and linked to a similarly modest increase in total metastatic burden (**Figure 4F**); burden at individual sites was mixed with no difference for bone metastases (**Figure 4G**). Bone-derived variants did not show restricted or enriched tropism or an enhanced ability to metastasize to bone when challenged by MIND or IC, but retained the ability to metastasize to multiple organs. However, IT injection highlighted that reintroducing these cells to the bone microenvironment greatly enhanced their metastatic potential. Bone-derived models grew rapidly after IT injection, with ∼10-fold greater whole-body BLI from ∼4 weeks post-injection onward (**Figure 4H**). IT challenge with the parental cells led to minimal metastatic burden within ∼14 weeks, whereas the bone-derived lines showed much greater total burden (**Figure 4I**), with several mice unexpectedly succumbing to metastatic disease. Per metastatic site, the parental cells showed minimal BLI consistent with background signal or micro-metastases only, whereas the bone-derived lines had extensive multi-organ macro-metastases, most notably to the uterus and ovaries (**Figure 4J**). Establishment of multi-route challenge models for the investigation of metastasis is important to capture the entire metastatic cascade. Bone derivative lines grew faster when challenged into bone, but their metastatic capabilities to bone were not pronounced when challenged by MIND or IC. Together, bone derived cell lines maintain multi-organ metastatic potential despite increased aggressiveness.

**Figure 4:**
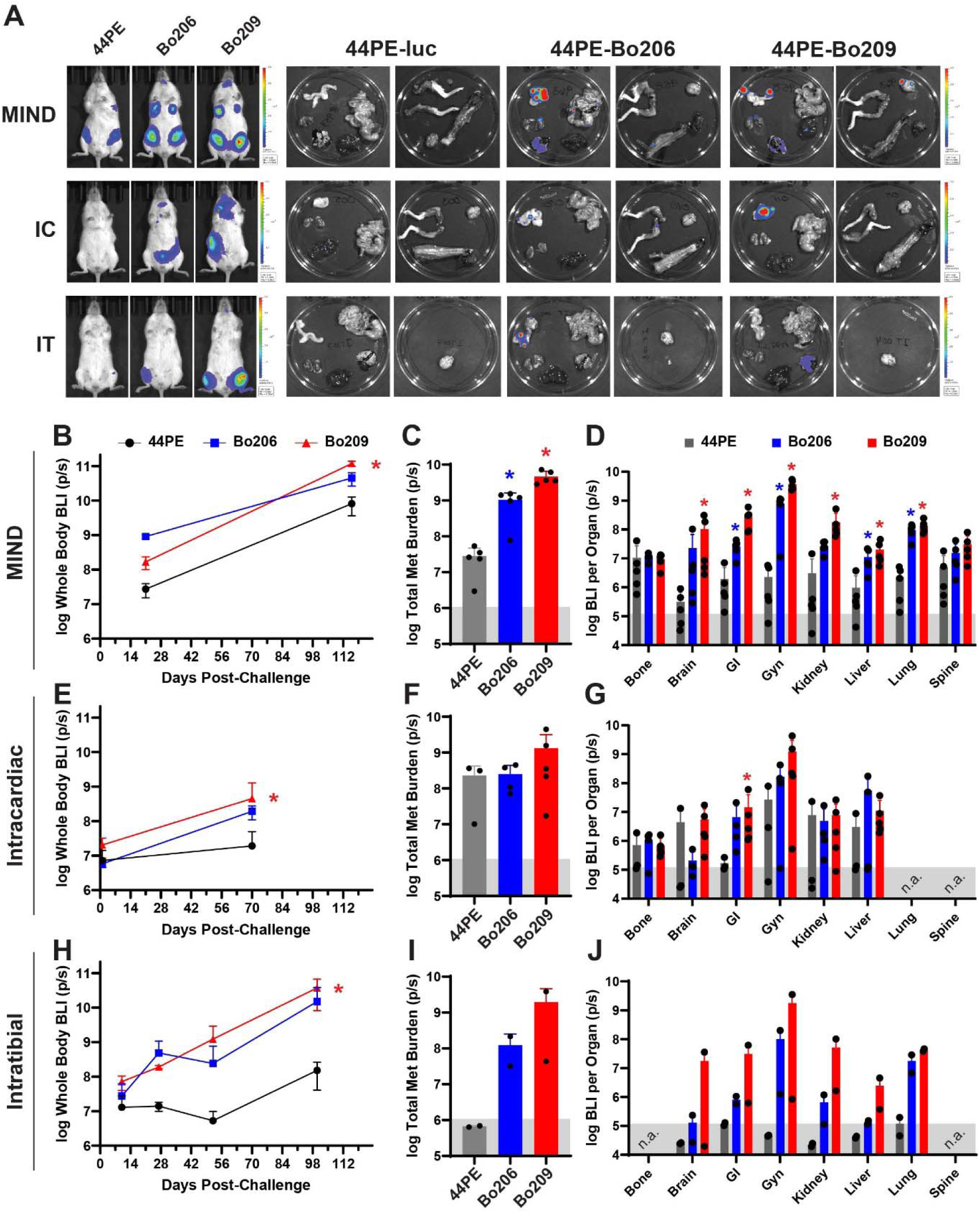
44PE bone derivative lines are more aggressive than the parental cell line. A) 44PE-luc (44PE), 44PE-Bo206 (Bo206), and 44PE-Bo209 (Bo209), were injected by MIND, intracardiac (IC), or intratibial (IT) routes into E2 supplemented mice (n=5/group). Representative bioluminescent images of whole body and *ex vivo* organs at time of humane endpoints. B/E/H) Whole body BLI measured over time; two-way repeated measures ANOVA. C/F/I) Total *ex vivo* metastatic burden (sum of all sites) measured by *ex vivo* BLI; one-way log-normal ANOVA. D/G/J) Metastatic burden as measured by *ex vivo* BLI of measured tissues; one-way log-normal ANOVA within each tissue. Significant (p<0.05) comparisons between 44PE-Bo206 and 44PE-luc (blue *) and 44PE-Bo209 and 44PE-luc (red *). Black bars depict two-way ANOVA adj.p<0.05. Background BLI signal (light-gray shaded boxes) determined by the average signal from a cohort of sham challenged animals.

### ILC maintains estrogen sensitivity in the bone microenvironment

Next, we examined the role for estrogen, and the potential shift in endocrine responsiveness, in the bone metastasis derived 44PE variants (**Figure 3**). First, we examined endocrine response of the derivative lines *in vitro*. In assessing proliferation, we noted that the bone-derived variants proliferated at a significantly reduced rate in culture versus parental 44PE-luc cells despite the aggressiveness of these cells *in vivo* (**Figure 4**).

However, 44PE-Bo206 and 44PE-Bo209 were responsive to the anti-estrogen fulvestrant, which suppressed cell growth (**Figure 5A**) and reduced the fraction of proliferative cells (**Figure 5B**), consistent with ER-driven growth. We also tested the converse by hormone-depriving cells in charcoal-stripped serum followed by treatment with E2, and similarly saw that parental and variant cells all ceased proliferation absent estrogen but were growth-induced on addition of E2 (**Figure 5C**). At the transcriptional level by RNA-seq, we found that ER-driven gene expression remained E2-responsive, as E2 treatment activated canonical ER targets as in the parental cells; 69% and 87% of genes in the Estrogene consensus signature [47] were still estrogen-regulated as in the 44PE parentals in Bo206 and Bo209, respectively (**Figure 5D**). These data support that the bone-derived variants remain estrogen-responsive and largely estrogen-dependent *in vitro*.

**Figure 5:**
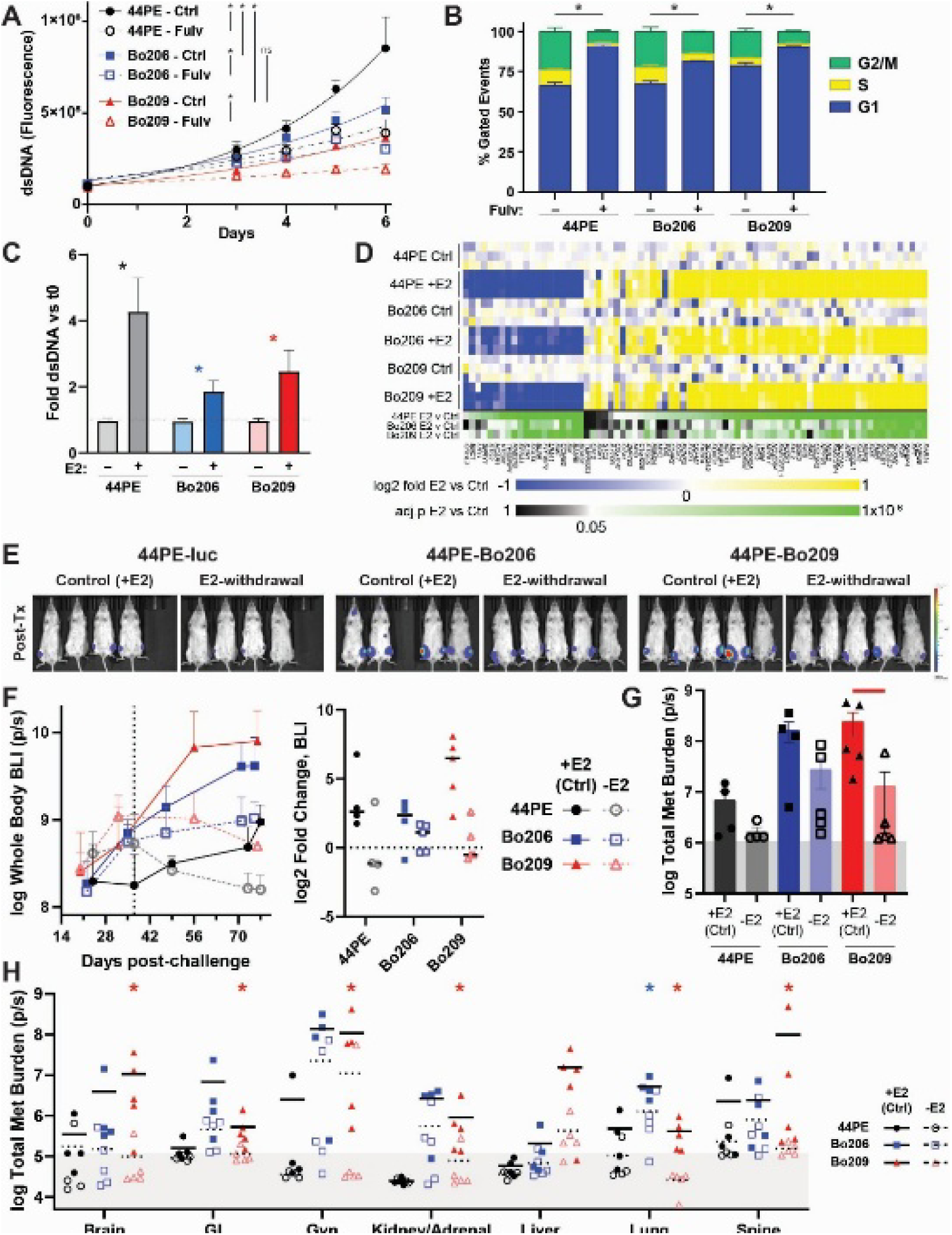
44PE bone derivatives maintain estrogen responsiveness but are less estrogen dependent. A) Proliferation assessed by dsDNA quantification after incubation in full serum media with and without Fulvestrant (100nM) for 6 days; significant comparison (* p<0.05) by one-way ANOVA with bar showing individual comparisons. B) PI based cell cycle analysis after 72 hours of fulvestrant (100nM) incubation; * p<0.05 by two-way ANOVA. C) Proliferation as measured by dsDNA quantification of cells hormone deprived then treated with E2 (100pM) for 6 days as normalized to day zero dsDNA concentration; * p<0.05 by t-test. D) Heatmap of canonical ER-targets from Estrogene depicting log2 fold change (blue-yellow) and adjusted p-value (p<0.05 green). E) Intratibial challenge of mice supplemented with 30µM E2 in the drinking water. After engraftment, E2 was withdrawn (-E2, to mimic aromatase inhibition) or maintained (+E2) for the duration of the experiment. E) Whole body BLI images of mice at humane endpoint. F) Quantification of whole-body BLI signal. G) Total metastatic burden as measured by *ex vivo* BLI; red bar represented p<0.05 by non-parametric t-test. Lung and spine were not measured due to residual post-injection disease in the thoracic cavity. H) Metastatic burden by organ as measured by ex vivo BLI. Significant (p<0.05) comparisons between 44PE-Bo206 and 44PE-luc (blue *) and 44PE-Bo209 and 44PE-luc (red *) determined by one-way lognormal ANOVA. Background BLI signal (light-gray shaded boxes) determined by the average signal from a cohort of sham challenged animals (n=10 animals; n=5 +E2 and n=5-E2).

The stark difference in *in vitro* vs *in vivo* growth rates of the bone-derived variants suggests important roles for both ER and the tissue microenvironment. To examine this, we engrafted 44PE-luc, 44PE-Bo206, and 44PE-Bo209 cells by IT injection in E2-supplemented mice, allowed tumors to establish for ∼5 weeks, then withdrew E2 and monitored progression versus control (+E2). As in **Figure 4**, the +E2 bone-derived models grew substantially faster than the parental cells (**Figure 5E-F**). E2-withdrawal reduced total body BLI in all three models modestly over ∼6 weeks of E2 withdrawal (**Figure 5E-F**), though statistical significance was not reached. Total *ex vivo* metastatic burden at endpoint was similarly reduced in 44PE-luc and 44PE-Bo206, and significantly reduced in Bo209 (**Figure 5G**), owing in part to greater total burden in control (+E2) mice. In evaluating metastatic burden per site (**Figure 5H**), metastases were limited in the parental cells within this timeframe, with the exception of spinal metastases. 44PE-Bo206 had a modest per site burden, with BLI in the lung significantly reduced by E2 withdrawal. 44PE-Bo209 had the greatest metastatic burden per site, which was significantly reduced by E2 withdrawal at nearly all sites. Notably, whereas liver metastases were again not observed with the parental 44PE-luc, both 44PE-Bo206, and to a greater extent 44PE-Bo209, presented with liver lesions.

Together these data highlight that *in vivo* metastatic adaptation renders ILC cells more aggressively metastatic, but this is not directly associated with an increased growth rate *ex vivo*, is critically dependent on the tissue microenvironment, and remains estrogen-driven. Notably, metastasis was not associated with specific organ tropism despite selection of these models from bone metastases, consistent with the shared gene expression changes see across models (**Figure 3**).

### ILC promotes a heterogenous yet pro-sclerotic phenotype in bone both in vivo and in vitro

mILC bone metastases have been commonly described as mixed osteosclerotic/osteolytic, in contrast to primarily lytic mNST bone lesions [7, 13-17, 48]. We examined the phenotype of bone lesions spontaneously forming after MIND challenge, and found that 44PE-luc tumors promote mixed osteosclerotic/osteolytic lesions in the spine (**Figure 6A**). Histologic analysis of a mouse challenged with 44PE-luc by MIND shows cortical breach by osteolytic disease in addition to immature osteoid deposition (**Figure 6B**). We then examined the phenotype in bone lesions after IT challenge, and observed a mixed osteosclerotic/lytic phenotype throughout the proximal tibia. Together, these data support that primary and metastatic bone disease in mice supplemented with E2 matches the clinical presentation of mILC to bone.

**Figure 6:**
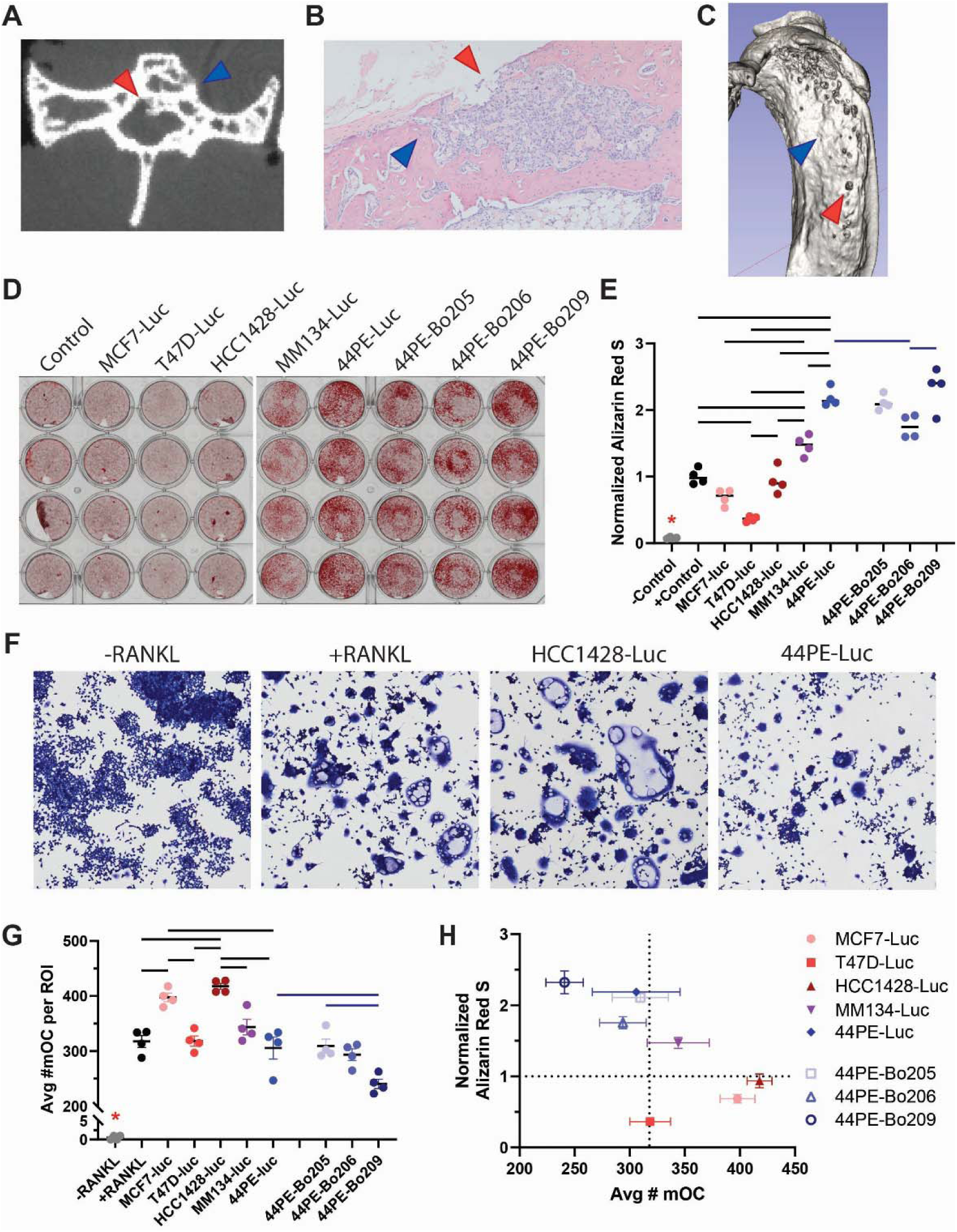
ILC bone metastases promote mixed lesions through activation of osteoblasts and inhibition of osteoclastogenesis. A) microCT of a lumbar vertebral metastasis developed from a 44PE-luc MIND challenged mouse with osteoblastic and osteolytic characteristics. B) Histologic (H&E) section of spontaneous tibial metastasis after MIND challenge of 44PE-luc. C) Representative microCT reconstruction of a mouse challenged with 44PE-luc show osteoblastic (blue arrow) and osteolytic (red arrow) changes. D) Conditioned media (CM) from NST (Red) and ILC cell lines (Blue/Purple) was cultured with MC3T3-E1 cells supplemented with ascorbic acid, β-glycerophosphate, and CaCl_2_ and stained with Alizarin Red S stain. E) Elution and quantitation of Alizarin Red S stain from wells shown in (A). CM from NST and ILC cell lines incubated with Raw264.7 cells, stained with Giemsa, and mature OC (mOC; 3+ nuclei per cell). F) Representative images of mOC after CM incubation. G) Quantification of the average number of mOC per 50x50pt (∼1.76cm x 1.76cm) ROI. H) Comparison of normalized Alizarin Red S stain (E) and number of mOC (G) shows ILC promote osteoblastic differentiation and mineral deposition and inhibited mOC differentiation, than NST lines. Black bars depict one-way ANOVA adj.p<0.05. Blue bars depict one-way ANOVA adj.p<0.05 comparison of bone derivative lines to the 44PE-luc parental cell line.

To investigate potential mechanisms driving the unique phenotype of mILC bone metastases, we investigated the effect of conditioned medium (CM) from ILC and NST cell lines on MC3T3-E1 (pre-osteoblast cells) to determine if factors secreted from ILC and NST cell lines have differential effects on osteoblast differentiation (the primary bone producing cells), matrix deposition, and mineralization as measured by Alizarin Red S staining. As reported previously, CM from NST cell lines inhibited mineral formation [49, 50] compared to control, showing limited induction of osteoblast differentiation. Conversely, CM from ILC lines strongly promoted mineral deposition (**Figure 6D-E**). We also examined osteoclast (the primary bone resorption cells) differentiation by incubating CM with Raw264.7 cells [34]. Consistent with their pro-osteolytic behavior, CM from NST cells induced extensive osteoclast differentiation and maturation while CM from ILC cells induced limited osteoclast differentiation; osteoclastogenesis was further reduced in CM from bone derived models (**Figure 6F-G**). Together, osteoblasts and osteoclasts regulate bone homeostasis, a process corrupted by tumor growth in bone. Comparting relative osteoblastic versus osteoclastic responses from these assays, CM from NST cells drove an overall pro-osteolytic phenotype, whereas CM from ILC cells drove a pro-osteoblastic phenotype (**Figure 6H**), likely reflective of the heterogenous bone lesions observed *in vivo*.

## DISCUSSION

There are few models that allow for the study of spontaneous ER+ mILC [29], therefore restricting understanding of the metastatic cascade and developing therapeutics specifically for mILC progression. Herein, we utilize the MIND model supplemented with estradiol in the drinking water to develop models of mILC which recapitulate the clinical presentation of disease including 1) the sites of metastatic dissemination (e.g. ovary and LMD), 2) maintenance of estrogen sensitivity, and 3) the unique interactions of mILC in distal organs (e.g. mixed osteosclerotic/osteolytic lesions in bone and LMD in CNS) without requiring resection of the primary tumor. Models of estrogen responsive metastatic disease will allow for therapeutic evaluation in the distinct context of mILC.

Metastatic organotropism is a complex process by which tumor cells may preferentially spread to organs supporting the unique phenotypes of the metastasizing cells, as first described in the seed-and-soil hypothesis [51]. The plasticity of both the seed and the soil has been widely debated with multiple studies showing restricted metastatic pathways to specific organs, pleiotropic, anatomic, and stochastic metastatic spread [52-54]. Many of these changes are considered transient, imprinted by the host tissue and exploited by a susceptible distal pre-metastatic niche. Interestingly, in our models, we observe multiple heterogenous bone derivative cell lines with increased primary tumor growth and metastatic potential, but do not preferentially re-seed the bone upon MIND or IC challenge. Increased growth observed upon IT challenge suggests that the selected cells prefer the bone microenvironment, but retain the capacity to metastasize and colonize other tissues. Bone has been associated with ER conversion and HR+HER2 induction leading to secondary metastatic development [55, 56]. With bone being the first site of metastasis diagnosed in a majority of mILC patients, it is feasible that bone acts as a reservoir and incubator for increased tumor aggressiveness; Whiteley et al showed that metastases between the bone and meninges are connected [57], and may partially explain these shared sites of metastasis in mILC. The metastatic behavior of ILC may be associated with E-cadherin loss leading to increased adaptability of the cells to multiple organ environments or the nutrient and growth-factor rich bone environment [43]. Notably, the hallmark E-cadherin loss of ILC has presented a quandary in metastatic progression since transient suppression and re-expression of E-cadherin is otherwise understood to be required for EMT, colonization, and outgrowth of metastatic disease [58]. With these advanced metastatic models, future work can dissect how ILC spread and colonize distinct metastatic niches.

Clinical diagnosis of ILC bone metastasis is complicated by the mixed osteosclerotic/osteolytic nature of the metastasis, coupled with the physiological context of bone loss associated with menopause, osteoporosis, and endocrine therapy. While bone scintigraphy and DEXA are important tools for identifying metastatic disease and bone mineral density, additional modalities are likely necessary to diagnose mILC. To date, PET imaging with fluorodeoxyglucose (FDG) and fluoroestradiol (FES) have been used monitor mILC, but questions remain concerning the types of lesions these radionuclides are best suited to [14, 15, 48, 59]. A recent investigation by Moustaquim et al showed FDG-PET/CT and FES-PET/CT had similar detection rates and noted the challenge in defining which radiotracer a patient may benefit from the most, suggesting that initial investigation be performed by FDG-PET/CT due to cost and accessibility [18]. In their study, the most common site of metastasis observed was bone. Recent evidence suggests that osteosclerotic metastases are non-FDG-avid compared to osteolytic lesions suggesting that FES-PET may be more effective in diagnosing mILC bone disease [15, 18, 48]. The other confounding factor is ER status of the metastatic disease which may lead to false-negative diagnoses by FES-PET [60]. ER expression has been associated with osteolytic disease, but activation of ER alone is insufficient to explain this phenotype in mILC [61, 62]. Pre-clinical investigation of bone metastatic phenotype in addition to *ESR1* expression, can be investigated in our MIND and IT metastasis models. In addition, the association between decreased FDG-avidity and osteosclerotic disease is poorly understood, but may benefit from comparison of heterogeneous metastases in a controlled investigative environment [63]. Identifying non-invasive methods of metastatic detection and monitoring are necessary due to the extended latent period of ILC recurrence and progression 5+ years after initial treatment cessation.

Bone is an estrogen responsive organ and the role of ER+ expression in bone maintenance has been well established [62]. MCF7 is a common ER+ osteolytic model of NST breast cancer bone metastasis; over-expression of the *neu* onocogene leads to an osteosclerotic phenotype in mice, without estrogen supplementation, by increased PDGF-BB expression [64, 65]. These data suggest an ER-independent mechanism leading to osteosclerotic metastases which may inform the mixed lesions associated with mILC. The factors and microenvironmental interactions leading the prominent mILC phenotype have yet to be defined.

Furthermore, exploitation of these variations for therapeutic gain has not yet been investigated. Current standard of care for breast cancer bone metastases are bisphosphonates and anti-RANKL therapies; however, these therapeutics have not been investigated directly in ILC patients [22, 66]. Recently, it has been suggested that anabolic therapy may be feasible in breast cancer patients [67]. Due to the osteosclerotic presentation of ILC bone metastases, evaluation of these therapeutics in ILC specific models is necessary, especially when combined with endocrine therapy which most patients will receive. The development of spontaneous metastatic models, with macro-metastatic symptomatic endpoints, will allow for therapeutic evaluation throughout the metastatic cascade, allowing for refinement of therapies specifically to patients with mILC.

## Authors’ contributions

JLS and MJS conceived the project and experiments. JLS and MJS designed and performed experiments and data analyses. JLS developed models for the project. All authors contributed to data analysis and interpretation. JLS and MJS wrote the draft manuscript; all authors read and revised the manuscript and have read and approved of this version of the manuscript.

## Funding

This work was supported by grants from the Cancer League of Colorado, Inc (JLS), the University of Colorado Cancer Center Cancer Research Training and Education Coordination (CRTEC) (JLS), Team Judy (JHO), the Lobular Breast Cancer Research Fund at CU Anschutz (MJS), support from the Women’s Cancer Developmental Therapeutics Program at CU Anschutz (MJS), and R01 CA251621 from the National Institutes of Health (MJS). This work utilized the U. Colorado Cancer Center Animal Imaging and Irradiation Shared Resource (RRID:SCR_021980), Genomics Shared Resource (RRID:SCR_021984), Biostatistics and Bioinformatics Shared Resource (RRID:SCR_021983), Cell Technologies Shared Resource (RRID: SCR_021982), and Flow Cytometry Shared Resource (RRID: SCR_022035) supported by P30 CA046934. The content of this manuscript is solely the responsibility of the authors and does not necessarily represent the official views of the National Institutes of Health or other agencies.

## Disclosures

None

## ACKNOWLEDGEMENTS

We would like to thank Dr. Jessica Finlay-Schultz and Dr. Carol Sartorius for providing us with the UCD4-GFP-luc cells. The authors appreciate the contribution to this research made by E. Erin Smith, HTL (ASCP)CM QIHC and Helen Dubois of the University of Colorado Cancer Center Pathology Shared Resource. This resource is supported in part by the Cancer Center Support Grant (P30CA046934). We thank Dr. Steffi Oesterreich for feedback on the manuscript.

## SUPPLEMENTAL FIGURE LEGENDS

**Supplemental Figure 1:**
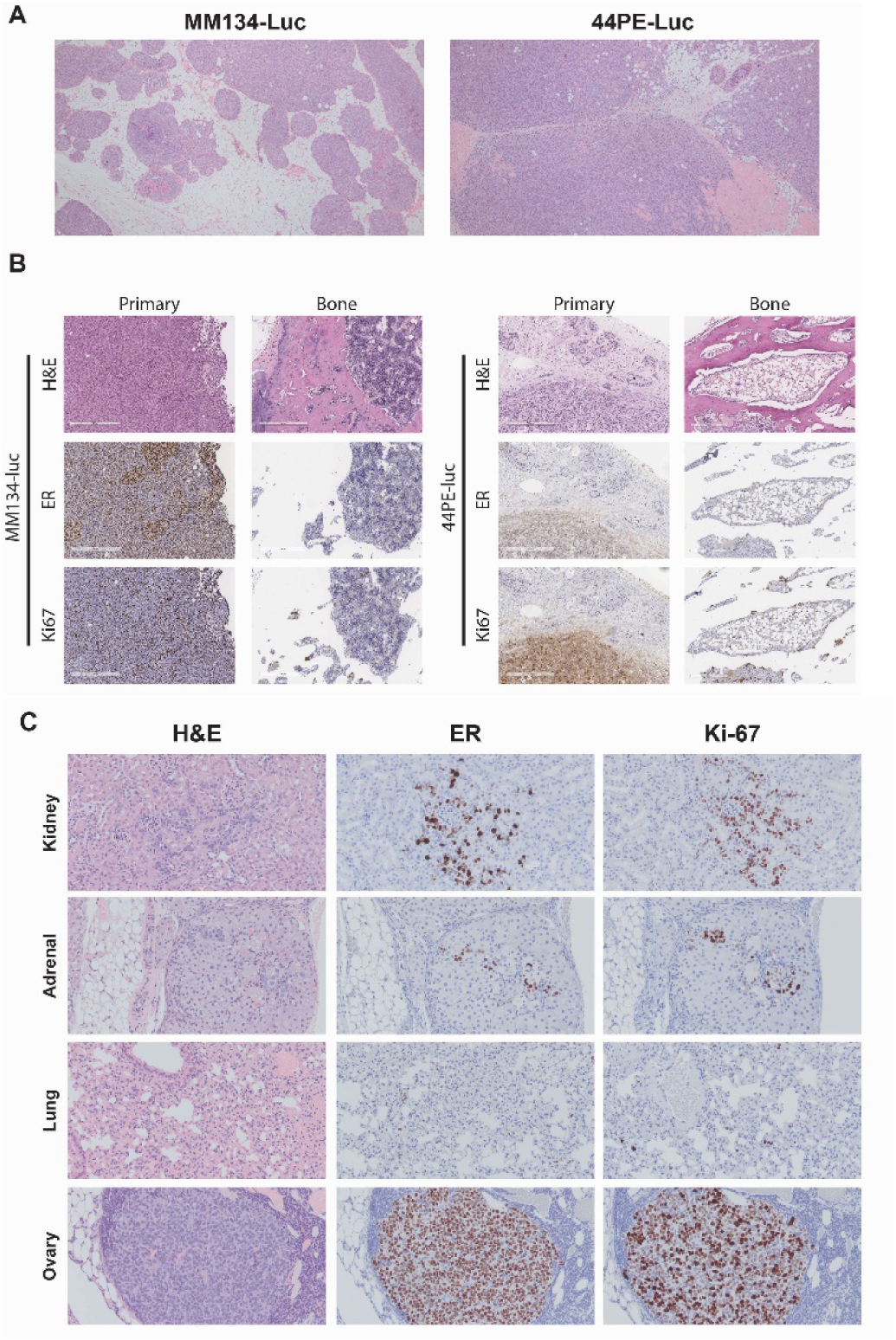
Histologic analysis of MIND tumors and resultant metastatic disease. A) MM134--Luc H&E MIND challenged tumor depicting an in situ like growth pattern. 44PE-Luc MIND challenged tumor depicting an invasive phenotype throughout the mammary gland. B) Histology of spontaneous metastatic development of MM134-luc and 44PE-luc from the primary tumor to bone stained with H&E, ER, and Ki67. Histology of MM134-luc spontaneous metastatic spread to kidney, adrenal gland, lung, and ovary analyzed by H&E, ER, and Ki-67 staining.

**Supplemental Figure 2.**
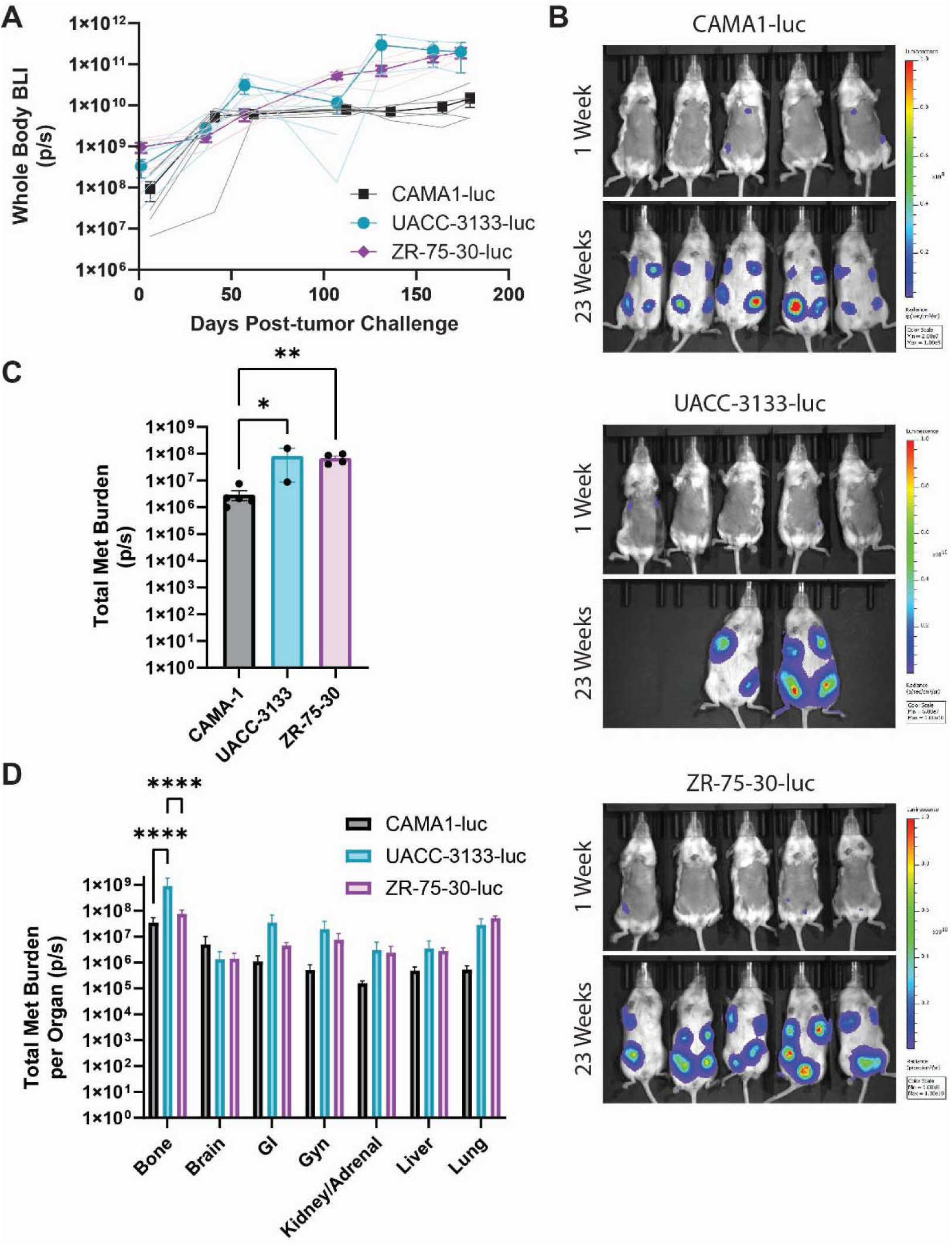
Metastatic potential of additional ILC lines. ILC lines CAMA1-luc, UACC-3133-luc, and ZR-75-30-luc were challenged via MIND into mice (n=5/cell line) supplemented with 30µM E2 in the drinking water. A) Whole body luminescence measured over the course of the experiment. Darker lines represent group averages until euthanasia of first mouse per group. Faded lines represent individual mice until their individual time of euthanasia. B) Representative images of mice at 1 week and 23 weeks after tumor challenge. Multiple UACC-3133-luc mice (n=3) succumbed to rapid metastatic progression leading to incomplete *ex vivo* BLI metastasis data. C) Total metastatic burden as measured by *ex vivo* BLI. One-way lognormal ANOVA: *adj.p<0.05; **, ANOVA adj.p<0.01. D) Metastatic burden per organ, as measured by *ex vivo* BLI. One-way lognormal ANOVA: *, ANOVA adj.p<0.05; **, ANOVA adj.p<0.01; ***, ANOVA adj.p<0.001; ****, ANOVA adj.p<0.0001.

**Supplemental Figure 3:**
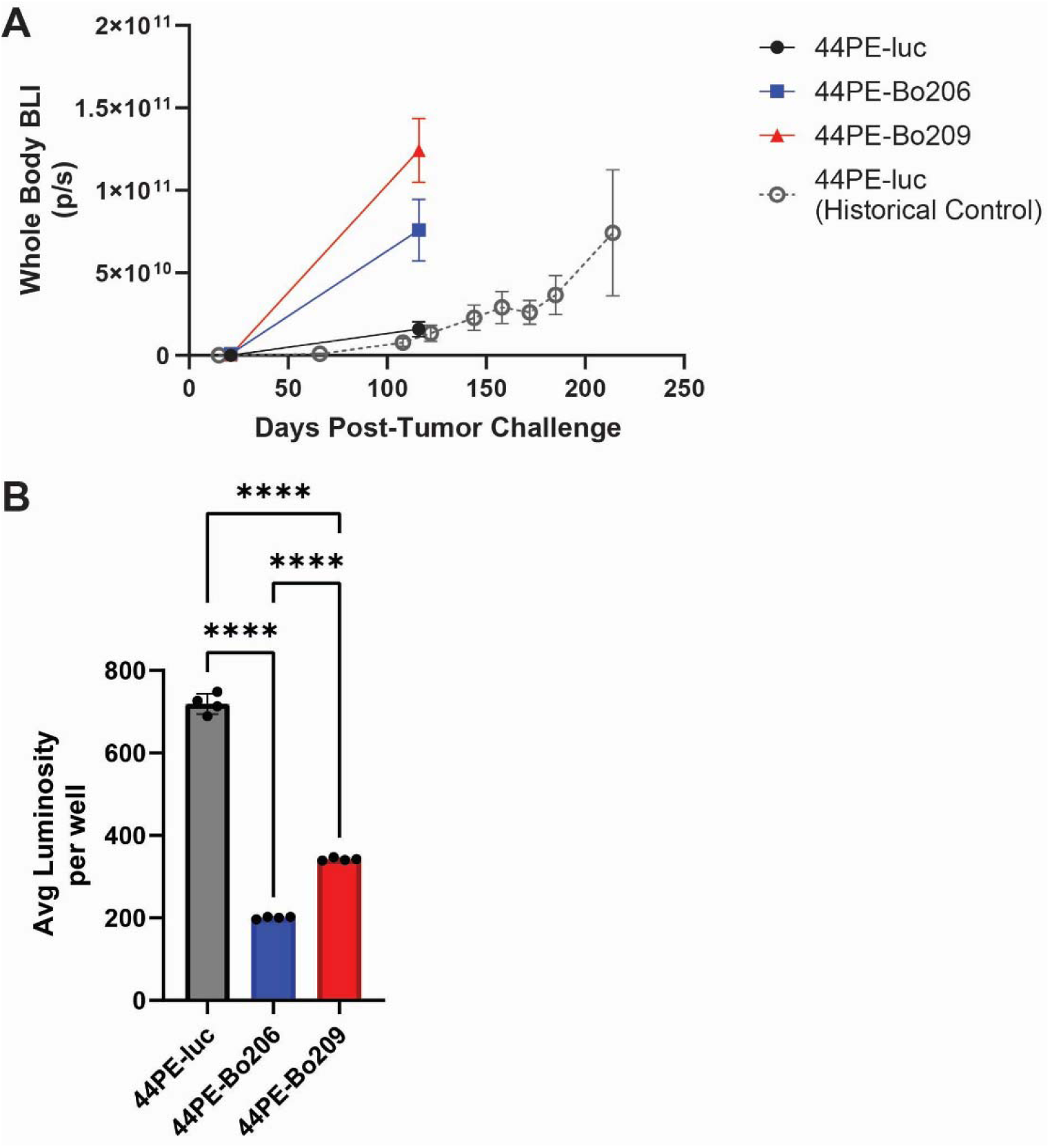
Growth and luminosity of 44PE bone derivative lines. A) MIND challenge of 44PE-luc and 44PE-Bo206/44PE-Bo209 bone derivative lines into E2 supplemented mice. Light gray line depicts historical growth after MIND challenge. While the current 44PE challenge is similar to historical studies, bone derivative lines grew substantially faster than expected. B) *In vitro* luminosity of cell lines. One-way ANOVA **** adj.p<0.0001.

**Supplemental Figure 4:**
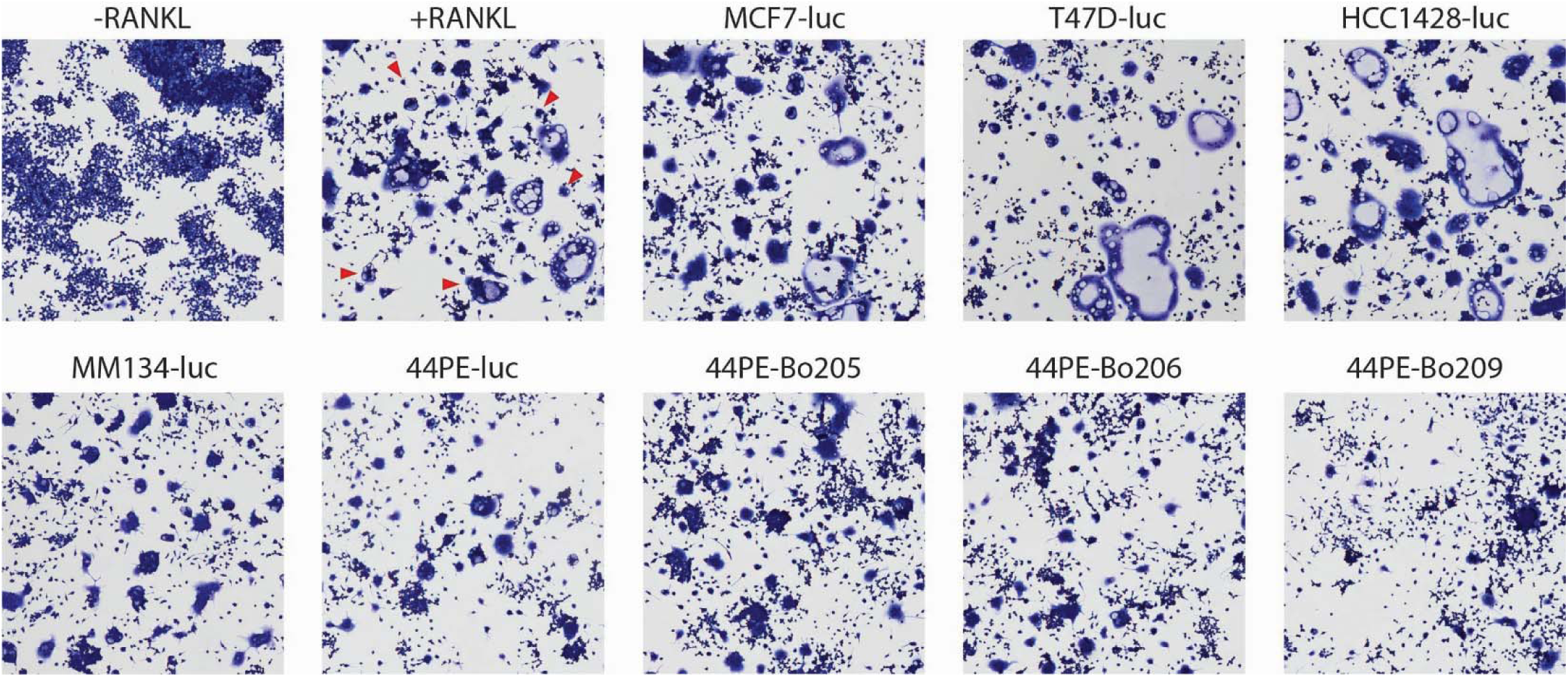
Bone derivative lines grow significantly faster than the parental line. CM was incubated with Raw264.7 cells and RANKL to induce differentiation into mOC (3+nuclei per cell). Representative images of May-Grunwald/Giemsa-Azur stained mOC are shown. The number of mOC were counted in 10 ROIs per well. Red arrows highlight mOC.

**Supplemental Table 1:**
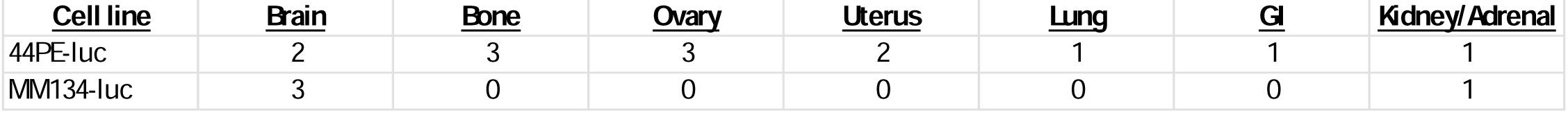
Established mILC organotropic cell lines.

## Notes

### Competing Interest Statement

The authors have declared no competing interest.

